# Copy number variation analysis of 9,482 *Mycobacterium tuberculosis* isolates identifies lineage-specific molecular determinants

**DOI:** 10.1101/2024.10.15.618387

**Authors:** Nikhil Bhalla, Anil Behera, Ashish Gupta, Ranjan Kumar Nanda

**Affiliations:** Translational Health Group, International Centre of Genetic Engineering and Biotechnology, New Delhi Component, Aruna Asif Ali Road, New Delhi, India

**Keywords:** CNV, Lineage, Virulence, AMR, Resistance, Tolerance

## Abstract

**Background:** Clinical manifestations of tuberculosis (TB) caused by *Mycobacterium tuberculosis* (Mtb) show lineage-specific differences contributed by genetic polymorphism such as phylo-single nucleotide variations (PhyloSNPs) and insertion or deletions (INDELs). Intragenomic rearrangement events, such as gene duplications and deletions, may cause gene copy number differences in Mtb, contributing to lineage-specific phenotypic variations, if any, which need better understanding.

**Results:** The relative gene copy number differences in high-quality publicly available whole genome sequencing datasets of 9,482 clinical Mtb isolates were determined by repurposing and modifying an RNA-seq data analysis pipeline. The pipeline included various steps, viz., alignment of reads, sorting by coordinate, GC bias correction, and variant stabilising transformation. The strategy showed maximum separation of lineage-specific clusters in two principal components, capturing ∼54% variability. Unsupervised hierarchical clustering of the top 100 genes and pairwise comparisons between Mtb lineages revealed an overlapping subset of genes (n=42) having significantly perturbed copy numbers (Benjamin Hochberg adjusted P-value < 0.05 and log_2_(drug-resistant/sensitive) > ± 1). These 42 genes formed multiple tandem gene clusters and are known to be involved in virulence, pathogenicity and defence response to invading phages. A separate comparison showed a significantly high copy number of phage genes and a recently reported druggable target Rv1525 in pre- and extensively drug-resistant (Pre-XDR, XDR) compared to drug-sensitive clinical Mtb isolates.

**Conclusion:** The identified gene sets in Mtb clinical isolates may be useful targets for lineage-specific therapeutics and diagnostics development.

## Background

A recent Global Tuberculosis report by the WHO showed an increased treatment success rate with a decreasing trend of mortality rates in Tuberculosis (TB) patients (1). Further, developing novel diagnostics and therapeutics might accelerate TB eradication efforts. Five lineages (lineage 1, 2, 3, 4 and 7) of *Mycobacterium tuberculosis* (Mtb), the pathogen responsible for TB, are adapted to infect humans (2). Isolated cases of zoonotic transmission of Mtb to humans have been reported, and specific lineages (such as lineage 5 and 6) are animal-adapted (3,4). Mtb has a unique strategy for developing drug resistance that includes the acquisition of mutations in primary or secondary drug target genes, which abrogates drug and drug-target interactions, leading to ineffective anti-mycobacterial activity. These mutations are commonly screened for drug resistance status determination and show high sensitivity and specificity comparable to *in vitro* culture-based drug susceptibility tests (DSTs) (5,6,7). Lineage-specific drug resistance-associated single nucleotide polymorphisms (SNPs) have been identified in various *rpoB* codons 450, 430, 435, 445, and 452 leading to amino acid substitutions. These substitutions impact Mtb’s *in vitro* growth kinetics, compromising culture-based phenotypic DST results (8,9). Association of specific clinical features and lineage of infecting Mtb strains, such as high frequency of pulmonary infection with lineage 2, 3 and 4; lung fibrosis with lineage 1; and bone TB disease with lineage 1, are well known. Lineage-specific differences in transmissibility are observed in Mtb. For example, Mtb of lineage 2 and 4 have high transmissibility compared to other lineages (10,11). In addition, lineage 2 is also known to have a high frequency of mutation acquisition, leading to higher drug resistance development. In contrast, lineage 4 is known to have high transmissibility but a low frequency of drug resistance development (12,13). Transcriptomic and proteomic profiling studies have also revealed subtle lineage-specific differences in overall biology (14,15,16).

Mtb genes of the H37Rv laboratory strain have been well-studied. However, genes and genome dynamics in clinical Mtb isolates need additional studies. The mycobacterial genome contains various genetic rearrangement-causing elements like transposases, insertion sequences and repetitive sequences (17,18). Insertion sequence like IS6110 is used as an epidemiological marker and reported to undergo transposition in the same strain in less than a year, indicating high transposition-like events in clinical Mtb isolates (18). A 350 kb genetic island duplication was reported in the laboratory Mtb strain (19). Some genetic regions are known to differ among lineages and are described as region of differences (RDs). These RDs are flanked by insertion sequences, which may show genetic rearrangement contributing to Mtb genome dynamics and affecting their phenotypes (20). These RDs aid in identifying Mtb lineages and molecular events caused by these genetic elements, which have probably played a major role in the evolution of present-day infectious clinical Mtb isolates. The copy number of *rrS* gene (16s rRNA) shows high variability in fast- and slow-growing mycobacteria. Its copy number directly correlates with growth rate as fast growers contain two copies of *rrS*, whereas a single copy is present in slow-growing mycobacteria (21,22,23). In addition, variations in the copies of insertion sequences, tRNAs, and ncRNAs genes are also well-known in Mtb (24,25,26,27). More than two copies of a set of Mtb genes (n=28) were reported in the genome sequences of various clinical Mtb isolates, highlighting the occurrence of events leading to gene copy number differences (28). A detailed study on the gene copy number differences in clinical Mtb isolates on a larger scale will provide a landscape of the genetic copy number variations in different clinical isolates.

In this study, we used publicly available whole genome sequencing raw data of ∼12,592 clinical Mtb isolates, repurposed, and optimized an RNA-seq data analysis pipeline to determine differences in the relative gene copy numbers in the clinical Mtb isolates belonging to multiple lineages with varying drug resistance profiles.

## Results

### High-quality Mtb WGS datasets (n=9,482) showed relative gene copy number differences

Il- lumina chemistry-based WGS data of 12,592 clinical Mtb isolates from 43 BioProjects were downloaded, and amongst all, 11,304 successfully underwent de novo assembly, and 11,234 underwent alignment, sorting and GC bias correction. A set of 1,125 *de novo* assemblies showed < 95% alignment to the Mtb reference genome, indicating contamination and were therefore excluded. A set of 295 samples could not be *de novo* assembled and thus had to be excluded. Out of these, 11,172 were successfully grouped based on their drug resistance and lineage status. Out of 11,172 clinical Mtb isolates, a subset (n=10,836) was from pure Mtb lineages, and the rest (n=335) showed mixed lineages. The lineage of one sample remained undetermined and was also excluded from subsequent analysis. Because of the low sample size, Lineage 5 (n=1), Lineage 6 (n=1), La1 (n=10) and La3 (n=22) were also excluded from subsequent analysis. Finally, 9,482 WGS datasets were found to be of optimal quality and selected for further processing (Figure 1A). Majority (n=6286; 62.9%) of the selected clinical Mtb WGS datasets were from India and the rest were from China (n=932; 9.3%), Pakistan (801; n=8%), Thailand (362; 1.7%), Israel (n=129; 1.3%), Myanmar (n=89; 0.9%), Cambodia (n=80; 0.8%), Oman (n=69; 0.7%), South Korea (n=58; 0.6%), Brazil (n=42, 0.4%), Iran (n=37; 0.4%), Lebanon (n=11; 0.1%), Malaysia (n=8; 0.1%) and mixed (n=575, 4.1%) (Figure 1B). Lineage and drug resistance-specific population distribution of the unfiltered (n=11,172) and filtered (n=9,482) Mtb WGS data are presented in Figures 1C and 1D.

**Figure 1:**
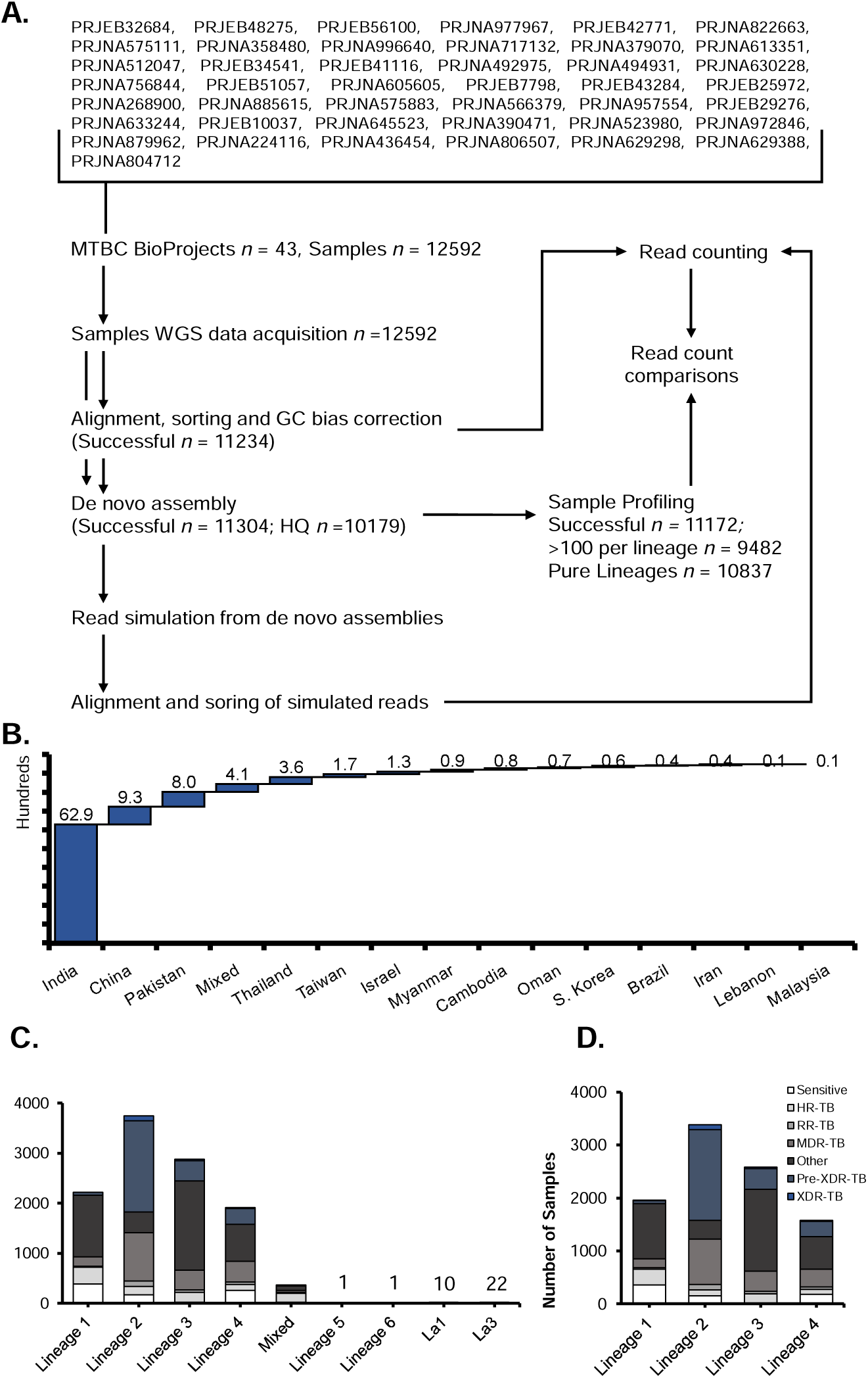
Data curation, processing and population structure of the whole genome sequence (WGS) data of clinical Mtb isolates used in this study. A: Schematic of data curation and processing steps followed. B: Mtb WGS data from the endemic countries and their distribution. C: The distribution of the WGS Mtb unfiltered data between lineages. D: Filtered (> 95% alignment with NC_000962.3 reference) WGS Mtb data distribution between major lineages. MTBC: *Mycobacterium tuberculosis* complex; HR: Isoniazid mono-resistant; RR: Rifampicin mono-resistant; Pre-XDR: pre-extensively drug-resistant; XDR-TB: extensively drug-resistant.

### GC bias correction followed by variant stabilizing transform showed maximum separation of lineage-specific clusters in principal component analysis

As the selected data sets were from different BioProjects and therefore had extreme variability, conventional tools commonly used for absolute copy number variant calling pipelines could not be implemented successfully. Therefore, an alternate strategy for relative gene copy number differences was optimized, akin to the RNA-seq analysis pipeline, with modifications to reduce the batch effects (Figure 2A). For this, various approaches were tested, all with minor modifications. Approach-I, which included alignment, sorting, and Variance Stabilizing Transformation (VST), showed diffused clusters of lineages on principal components that captured 24.5% and 15.3% variability. Approach-II included all the steps in Approach-I and GC bias correction of bam files before read count matrix preparation and VST. It showed lineage-specific cluster separation on principal components that captured 44.4% and 10.9% variability. Approach III was the same as the second, except that it included a batch correction step with the Combat tool after VST and showed similar patterns on PCA on principal components that captured 40.2 and 12% variability. The fourth approach included *de novo* assembly and read simulation using assemblies, followed by alignment and sorting. This strategy was impacted by outliers in the sample set but seemed to cluster samples according to lineages on principal components that captured only 21.2% and 15.6% variability (Figure 2B). The second and third approaches showed maximum separation of clusters with high variability in their principal components. Combat-assisted batch correction seemed unnecessary as without this step, the lineage-specific clustering of samples was apparent on the PCA that captured >54 variability, and thus, Approach-II was deemed an optimized method for relative gene copy number determination.

**Figure 2:**
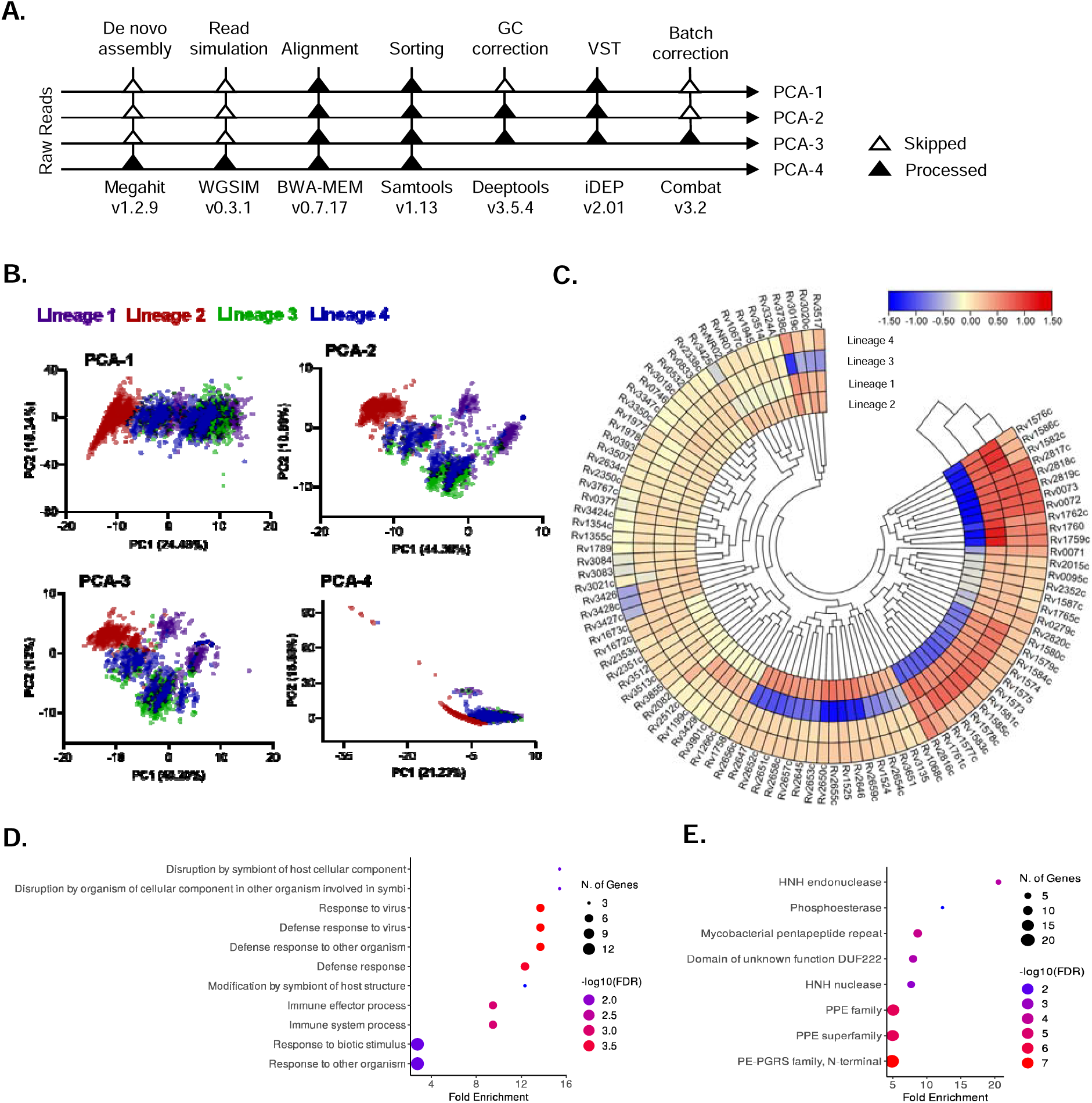
Data analysis optimization and multi-group analysis. A: The WGS data of clinical Mtb isolates was processed using four strategies to estimate gene copy number differences. The first approach included alignment, sorting and VST; the second approach included the same steps as well as GC correction before VST; the third strategy included batch effect correction using combat and VST; and the fourth strategy included de novo assembly, read simulation, alignment and sorting. The read count matrix was assessed using PCA analysis on the iDEP platform. B: PCA plots showing the lineage-specific distributions observed from the read count matrix prepared using the four approaches. C: The top 100 genes were subjected to hierarchical clustering. The Z-score medians of the top 100 genes per lineage were calculated and represented. D: Gene enrichment analysis of the top 200 genes determined after hierarchical clustering using the GO database. E: Protein domain enrichment analysis of top 200 genes after hierarchical clustering using InterPro database. PCA: Principal Component Analysis; PC1/2: Principal Component 1 or 2.

### Multi-group analysis revealed perturbations in the copy number of genes involved in bacterial immune response and associated pathways

Unsupervised hierarchical clustering of the top 100 genes after VST is shown in Supplementary Figure 1. The averaged VST normalized data of all samples in each lineage showed gene copy number perturbations in various tandem genes with respect to Mtb H37Rv reference genome viz. Rv0071-73, Rv1067c-68c, Rv1354c-55c, Rv1524-25, Rv1573-87c, Rv1672c-73c, Rv1758-62c, Rv1977-78, Rv2350c-53c, Rv2645-47, Rv2650c-59c, Rv2816c-20c, Rv3018c-21c, Rv3019c-20c, Rv3083-84, Rv3424c-29, Rv3512-1 and RvNR01-02. PE and PPE genes, Rv1355c (*moeY*), Rv3084 (*lipR*), Rv2351c (*plcA*) and Rv3855 (*ethR*) are non-tandem and were also found to have perturbed gene copy number differences between lineages (Figure 2C). Gene ontology clustering showed enrichment of disruption by symbiont of host cellular component (Rv2349c, Rv2350c, Rv2351c); disruption by an organism of a cellular component in other organism involved in symbiotic interaction (Rv2349c, Rv2350c, Rv2351c); response to the virus (Rv2816c-21c); defense response to the virus (Rv2816c-21c); defense response to other organisms (Rv2816c-21c); defense response (Rv2816c-21c); modification by symbiont of host structure (Rv2349c-51c); immune effector process (Rv2816c-21c); immune system process (Rv2816c-21c); response to biotic stimulus (Rv0404, Rv1266c, Rv1506c, Rv1917c, Rv2082, Rv2816c-21c, Rv3343c, Rv3377c-78c) and response to other organisms (Rv0404, Rv1266c, Rv1506c, Rv1917c, Rv2082, Rv2816c-21c, Rv3343c, Rv3377c-78c) (Figure 2D). Similarly, protein domain enrichment using Interpro database showed enrichment of HNH endonuclease (Rv1148c, Rv1765c, Rv1945, Rv2015c); Phosphoesterase (Rv2349c-51c); Mycobacterial pentapeptide repeat (Rv0304c-05c, Rv0355c, Rv1917c, Rv2353c, Rv3343c, Rv3347c, Rv3350c); Domain of unknown function DUF222 (Rv0095c, Rv0393, Rv1148c, Rv1587c, Rv1765c, Rv1945, Rv2015c); HNH nuclease (Rv0393, Rv1148c, Rv1587c, Rv1765c, Rv1945, Rv2015c); PPE family (Rv0305c, Rv0355c, Rv1789, Rv1917c, Rv2352c, Rv3018c, Rv3022c, Rv3135, Rv3343c, Rv3347c, Rv3350c, Rv3425, Rv3426, Rv3429, Rv3738c, Rv3739c); PPE superfamily (Rv0305c, Rv0355c, Rv1789, Rv1917c, Rv2352c, Rv3018c, Rv3022c, Rv3135, Rv3343c, Rv3347c, Rv3350c, Rv3425, Rv3426, Rv3429, Rv3738c, Rv3739c) and PE-PGRS family, N-terminal (Rv0109, Rv0279c, Rv0532, Rv0746, Rv0747, Rv1067c, Rv1068c, Rv1243c, Rv1452c, Rv1759c, Rv1768, Rv1788, Rv1803c, Rv2162c, Rv2490c, Rv2634c, Rv3388, Rv3507, Rv3508, Rv3511, Rv3595c) (Figure 2E).

The top 100 genes could also be categorized according to their functions relevant to understanding various aspects of virulence and the emergence of drug resistance. These categories included the following categories.

a) PE-PGRS and PPE-family: Rv0279c (PE_PGRS4), Rv0532 (PE_PGRS6), Rv0746 (PE_PGRS9), Rv0833 (PE_PGRS13), Rv1067c (PE_PGRS19), Rv1068c (PE_PGRS20), Rv1759c (Wag22), Rv3507 (PE_PGRS53), Rv3512 (PE_PGRS56), Rv3514 (PE_PGRS57), (Rv1789 (PPE family protein PPE26), Rv2352c (PPE38), Rv2353c (PPE39), Rv3018c such as PPE46), Rv3135 (PPE50), Rv3347c (PPE55), Rv3350c (PPPE56), Rv3425 (PPE57), Rv3426 (PPE58), Rv3429 (PPE59).
b) CRISPR-associated genes: Rv2816c-20c.
c) Antibiotic resistance genes: Rv1266c (serine/threonine-protein kinase PknH), Rv3855 (HTH-type transcriptional repressor EthR), Rv1355c (molybdopterin biosynthesis protein MoeY).
d) DNA-modifying enzymes: Rv1199c (IS1081 transposase), Rv2646 (Integrase), Rv3427c-28c (Transposases).
e) ESAT-6-like secretory protein: Rv3019c-20c (*esxR-esxS*).
f) Ribosomal RNA genes: RvNR01-02, 23S rRNA (*rrL*) and 16S rRNA (*rrS*).

### Pairwise comparisons between lineages and subsequent post hoc tests revealed a gene set identified via multi-group analysis

To determine the relative gene copy number differences, the DeSeq2 R package was repurposed. With respect to lineage-1, in lineage 2 and −3, relatively high copy numbers of Rv1524, Rv1525, Rv2645, Rv2646, Rv2647, Rv2650c, Rv2651c, Rv2652c, Rv2653c, Rv2654c, Rv2655c, Rv2656c, Rv2657c, Rv2658c, Rv2659c, Rv3135, and Rv3651 genes were observed. With respect to other lineages (1, 3 and 4), lineage 2 had relatively low gene copy numbers of Rv0072, Rv0073, Rv1573-86c, Rv1759c-62c, Rv2816c-19c. A low relative copy number of Rv0071 was observed in lineage 2 compared to lineage 4. Lineage 1, with respect to lineage 4, showed relatively low gene copy numbers of Rv15-25, Rv2645, Rv2646-47, Rv2650c-59c, Rv3135, and Rv3651 (Figure 3A). Unsupervised clustering of averaged lineage-specific normalized read counts of these genes also showed gene copy number differences in tandem genes viz., Rv2045-59c; Rv1759c-62c; Rv1581c-86c; Rv1573-75; Rv1576c-85c; Rv2816c-19c and Rv0071-72 matching with the results from multigroup analysis as described above.

**Figure 3:**
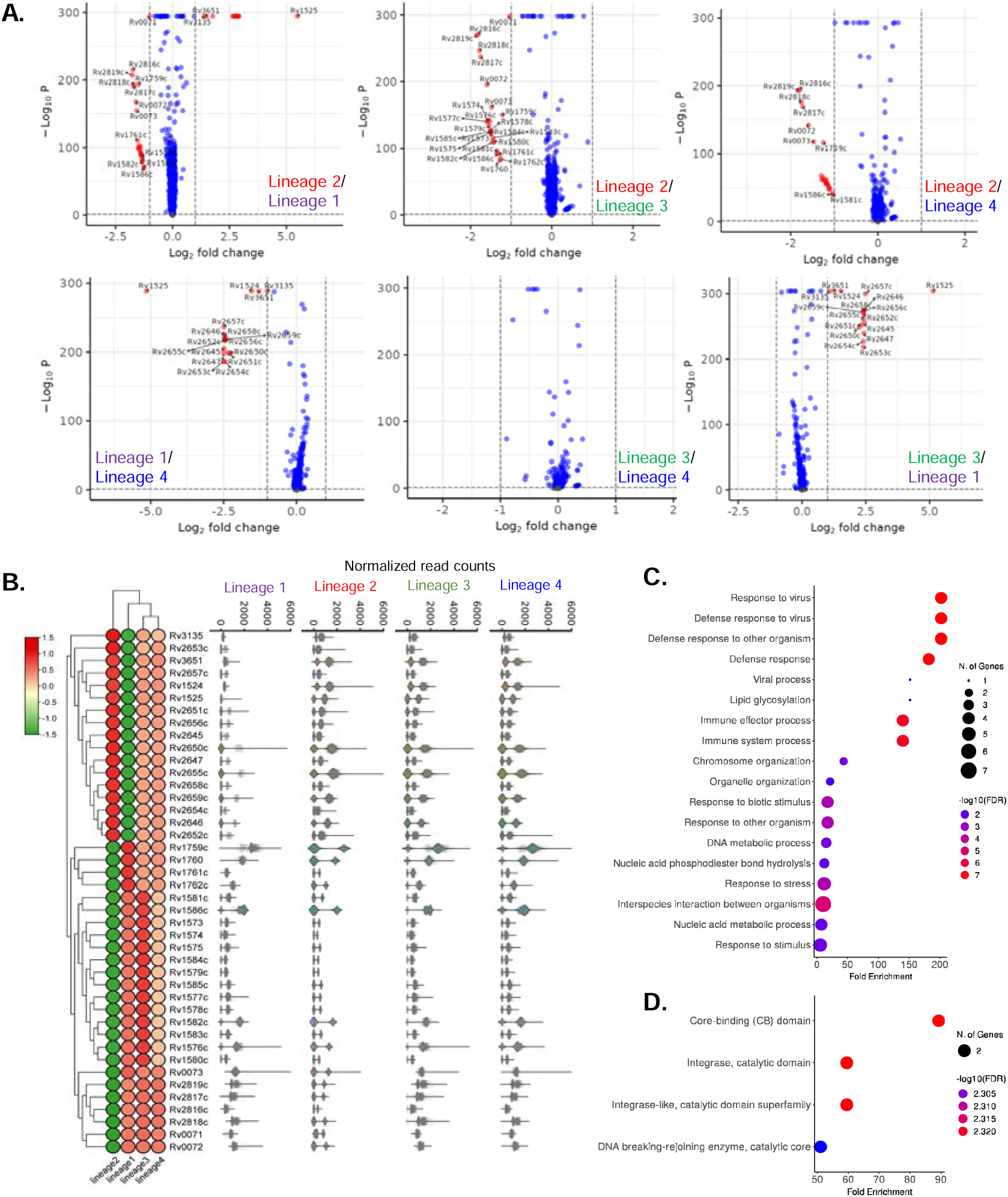
Lineage-specific gene copy number differences and effect sizes analysis showed sub-populations in different lineages. Normalization and statistical modelling were performed using DESeq2’s built-in functions. Pairwise comparisons between all experimental conditions were carried out, and the P-values were adjusted for multiple testing using the Benjamini-Hochberg method. A: The Log_2_(Read count FC) was plotted against –Log_10_(BH P-value) to represent the data in volcano plots. B: A set of 42 genes had differential gene copy numbers between lineages. The normalized read counts were averaged for each of these genes in all lineages and plotted as a heatmap to give an overview of gene copy numbers in Mtb lineages. To further get insights on lineage sub-populations with high or low gene copy numbers, the normalized read counts were plotted as violin plots, which showed subpopulations within each lineage with high and low gene copy numbers. However, a subpopulation is missing in specific lineages. C: Gene ontology clustering of the deregulated 42 gene set (GO database) showed perturbed pathways. D: Protein domain enrichment analysis (Interpro database) of these 42 deregulated genes showed varied protein domains. P: P-value, FDR: False Discovery Rate.

Lineage 2 had high copy numbers of Rv3135, Rv2653c, Rv3651, Rv2657c, Rv1524-25, Rv2651c-52c, Rv2656c, Rv2650c, Rv2645-47, Rv2654c-55c, and Rv2658c-59c genes compared to lineage 1. These gene sets had medium gene relative copy numbers in lineage 3 and lineage 4, whereas lineage 1 had the least copy number. Rv1759c-62c genes had high copy numbers in lineage 1, medium in lineage 3 and 4 and least in lineage 2. Rv1580c-84c, Rv1573-75, Rv1585c-86c, Rv1577c, and Rv1576c-79c genes showed maximum gene copy number in lineage 3, medium copy number in lineage 1 and then lineage 4 and least in lineage 2. Genes like Rv2816c-19c and Rv0071-73 showed high gene copy numbers in lineage 1, 3, and 4 and low gene copy numbers in lineage 2. Upon further analysis of normalized read counts, we found subpopulations of Mtb lineages that had either high or low copy numbers of specific genes within each lineage. In other words, the analysis showed a population that showed close to zero normalized read counts, indicating gene absence or low relative gene copy number in one subpopulation, and a second subpopulation having higher read counts, indicating gene presence and potentially high gene copy number (Figure 3B).

Gene ontology clustering showed enrichment of pathways like response to virus (Rv2816c-19c); defense response to virus (Rv2816c-19c); defense response to other organism (Rv2816c-19c); defense response (Rv2816c-19c); viral process (Rv2659c); lipid glycosylation (Rv1524); immune effector process (Rv2816c-19c); immune system process (Rv2816c-19c); chromosome organization (Rv2816c-17c); organelle organization (Rv2816c-17c); response to biotic stimulus (Rv2816c-19c); response to other organisms (Rv2816c-19c); DNA metabolic process (Rv2659c, Rv2816c-17c); nucleic acid phosphodiester bond hydrolysis (Rv2816c-18c); stress response (Rv1760, Rv2816c-19c); interspecies interaction between organisms (Rv0072, Rv1760, Rv2659c, Rv2816c-19c); nucleic acid metabolic process (Rv2659c, Rv2816c-18c) and response to stimulus (Rv1760, Rv2816c-19c) (Figure 3C). Protein domain enrichment showed enrichment of Core-binding (CB) domain; Integrase, catalytic domain; Integrase-like, catalytic domain superfamily; DNA breaking-rejoining enzyme, catalytic core with all these domains being contributed by only two genes: Rv2646, Rv2659c (Figure 3D).

### Pairwise comparison revealed a high copy number of Phage genes in drug-resistant clinical Mtb isolates

From the different drug resistance categories, like Rifampicin-, Isoniazid-resistant (RR, HR), multidrug-resistant (MDR), pre-extensively drug-resistant (preXDR) and XDR, the preXDR and XDR groups showed significant differences in the relative gene copy numbers compared with drug-sensitive Mtb clinical isolates (Table 1). These included Rv1525, Rv2654c, and Rv2657c-58c genes that showed a high copy number in the preXDR group, whereas the majority of the genes belonging to the Rv2846-59c cluster were higher in the XDR group, and the rest were similar.

**Table 1:** Copy number differences in genes of drug-resistant and sensitive clinical Mtb isolates. The DeSeq2 determined differential copy numbers of genes having Log_2_FC>±1.0 with BH adjusted P-value < 0.05 were considered significant. Only significant gene copy number differences between PreXDR/Sensitive and XDR/Sensitive were presented. XDR: Extensively drug-resistant; TB: Tuberculosis; BH: Benjamin Hochberg; FC: Fold Change.

| S. No. | Comparison | Gene | Locus tag | $\text{Log}_2\text{FC}$ | BH adjusted P-value |
| --- | --- | --- | --- | --- | --- |
| 1 | Pre XDR-TB vs Sensitive | Rhamnosyl transferase WbbL | Rv1525 | 1.2 | 2.50E-28 |
| 2 |  | Antitoxin | Rv2654c | 1.0 | 1.24E-18 |
| 3 |  | Prophage protein | Rv2657c | 1.0 | 1.29E-22 |
| 4 |  |  | Rv2658c | 1.0 | 1.02E-21 |
| 5 | XDR-TB vs Sensitive | Rhamnosyl transferase WbbL | Rv1525 | 1.3 | 1.45E-07 |
| 6 |  | Hypothetical protein | Rv2645 | 1.1 | 2.28E-05 |
| 7 |  |  | Rv2647 | 1.1 | 5.68E-05 |
| 8 |  | Integrase | Rv2646 | 1.1 | 4.64E-06 |
| 9 |  | Toxin | Rv2653c | 1.1 | 0.0001 |
| 10 |  | Antitoxin | Rv2654c | 1.2 | 1.16E-05 |
| 11 |  | Prophage protease | Rv2651c | 1.1 | 5.23E-06 |
| 12 |  | Prophage integrase | Rv2659c | 1.1 | 6.82E-06 |
| 13 |  | Prophage protein | Rv2650c | 1.1 | 1.05E-05 |
| 14 |  |  | Rv2652c | 1.2 | 4.89E-06 |
| 15 |  |  | Rv2655c | 1.1 | 1.32E-05 |
| 16 |  |  | Rv2656c | 1.1 | 8.32E-06 |
| 17 |  |  | Rv2657c | 1.2 | 2.43E-06 |
| 18 |  |  | Rv2658c | 1.2 | 2.64E-06 |

## Discussion

Clinical Mtb isolates are classified into lineages depending on their PhyloSNP pattern, known as barcodes or the presence or absence of large deletions occurring in RDs (17,20,29). The Mtb genome contains tandem and non-tandem genetic elements like repetitive DNA sequences such as IS6110 and direct short and long repetitive stretches. In addition, its genome also encodes proteins like integrases, resolvases, and transposases that predispose the Mtb genome to intragenomic recombination events that may lead to gene deletions and duplications (18). A report on pathogenic *Staphylococci* and *Enterococci* showed duplication of *csa1A*, *deoC*, *hsdM*, *rpmG*, *splF*, *bsaA2* and *vraH* genes and *glsB*, *lysM*, PTS lactose and transporter subunits, respectively (30). An existing report demonstrated a massive gene duplication of the Rv3128c-Rv3427c segment that encompasses 8% of the Mtb genome and 300 genes in clinical isolates (19). Isolated studies on the impact of copy number variations in different organisms, including prokaryotes and eukaryotes, indicate that the gene copy number differences correlate highly with expression profiles (31,32). These factors formed the basis of the present study, aiming to identify relative gene copy number differences that may cause known lineage-specific phenotypic differences and clinical manifestation changes amongst prevalent and circulating major Mtb lineages.

Conventional copy number variant calling tools such as CNVKit and CNVnator could not be successfully implemented due to large sample size and sample heterogeneity, so the existing RNA-seq data analysis pipeline was repurposed with modifications to determine relative gene copy number differences. GC bias correction is critical in copy number variant calling pipelines and thus, it was incorporated before the read-counting step approach in our optimized approach (33,34). The optimized approach yielded maximum separation of lineage-specific clusters on principal components that captured the highest variability. The repurposed RNA-seq pipeline with modifications could effectively be used for relative gene copy number analysis. From the hierarchical clustering analysis, the selected top 100 genes, had perturbed gene copy number differences amongst Mtb lineages. These genes were functionally categorized as PE-PGRS and PPE-family genes known to be involved in virulence and pathogenicity, with details highlighted in the results section. Lineage-specific virulence and pathogenicity changes are well known, and our findings on the lineage-specific copy number differences of these virulence-associated genes corroborate with existing literature (13,17,35). Amongst the identified PE and PPE genes, Rv3347c, Rv3347c and Rv3350c are known constituents of an RD known as RDrio, and Rv3738 constitutes N-RD25das and N-RD25_tbA. Therefore, their gene copy differences in clinical Mtb isolates could be attributed to the known general molecular behaviour of RDs (20). Also, the repetitive nature of these genes and their effect on the immune response post-infection and extensive involvement in host-pathogen interactions is described in earlier reports (35).

CRISPR-associated genes (Rv2816c-20c), which also form a portion of RD207 were identified by the multigroup, pairwise and post hoc analysis in this study (20). These genes are involved in DNA integrity, and their absence in lineage 2 and drug-resistant clinical Mtb isolates is known and corroborates our present findings that their gene copy number is low in lineage 2 compared to the rest (36,37). Antibiotic resistance-conferring genes Rv1266c, Rv3855, and Rv1355c, constitute RD145 (20). A higher gene copy number of Rv1266c was observed in lineage 3 compared to the rest of the lineages. Mutations in Rv1266c are associated with Delamanid resistance and are absent in drug-resistant clinical Mtb isolates (36). A higher copy number of Rv1266c in lineage 2 clinical Mtb isolates indicates an alternate mechanism of Delamanid resistance development apart from the acquisition of mutations. Rv3855 showed a relatively lower gene copy number, possibly contributing to developing ethionamide-specific drug tolerance. DNA-modifying enzymes like Rv1199c (IS1081 transposase), Rv2646 (Integrase), Rv3427c-28c (Transposases) showed a relatively higher gene copy number in lineage 1 than in other lineages. Reports indicating their differential abundance in Mtb lineages are limited, and these observations seem highly significant. Lineage-specific differences in dormancy induction and resuscitation and lower expression of Rv3427c-28c are previously known. Therefore, its perturbed copy number in Mtb lineages may have a role in some aspects of dormancy (38,39). ESAT-6-like secretory protein-encoding genes Rv3019c-20c (esxR-esxS) are previously known to duplicate and give rise to PE and PPE family virulence and host-pathogen interactome genes (17). ESAT-6 genes Rv3019c-20c had perturbed gene copy numbers amongst lineages, indicating their ongoing evolutionary duplication events in clinical Mtb isolates. Ribosomal RNA genes (RvNR01-02, 23S and 16S rRNA) copy number differences are previously known, viz., one copy and two copies of 16s rRNA in slow and fast-growing *Mycobacteria*, respectively. 16S rRNA gene has a high copy number in several non-mycobacterial species, suggesting its predisposition to duplication (27,40). Phage-related genes (Rv1573-86c) were recently annotated and were previously known as conserved hypothetical genes. We have observed differences in their gene copies in a lineage-specific manner, and limited literature details are available; therefore, exploring the role of these genes in host-pathogen interactions could be useful.

Rv1525 gene (rhamnosyl transferase WbbL) showed a high copy number in lineage 2, low or absent in lineage 1, and medium in lineages 3 and 4. It had a higher gene copy number in drug-resistant clinical Mtb isolates than drug-sensitive ones. Rv1525 has recently been described as a novel druggable target, and its duplication may contribute to drug resistance development upon exposure to its inhibitors (41). Domenech et al. reported duplication involving 300 Mtb genes, including Rv3128c to Rv3427v, in clinical Mtb isolates. They attributed the duplication of the Rv3128c-Rv3427c cassette to insertion sequences present upstream of Rv3128c and downstream of Rv3427c. The present study observed copy number differences in Rv3135, Rv3324A, and Rv3347c. Rv3350c, Rv3424c, Rv3425, Rv3426, and Rv3427c Mtb genes corroborate a previous report on Rv3128c-Rv3427c cassette duplication (19).

The pairwise and subsequent post hoc analysis led to the identification of 42 genes that showed differences in copy numbers amongst the lineages. These genes were a subset of the top 100 genes determined through multigroup analysis. It revealed subpopulations of lineages with high and low copy numbers, suggesting that duplicated genes have different resolving powers if they are to be used for delineating Mtb lineages. However, combining these genes in a CNV-based diagnostic test may provide high sensitivity and specificity in delineating Mtb lineages.

## Conclusion

The alternative strategy of determining gene copy number differences by repurposing conventional RNA-seq data analysis pipelines is possible with minor modifications by including the GC bias correction. The genes with perturbed relative copy number differences amongst Mtb lineages may form the basis of future diagnostics.

## Methodology

### Data acquisition and processing

The BioProject IDs were text-mined in conventional literature using databases like PubMed and Google Scholar from studies carried out primarily in India and other TB-endemic countries (Supplementary Table 1). The WGS data of reported clinical Mtb isolates from TB-endemic countries was acquired from the European Nucleotide Archive through the Wget script. The WGS data in fastq format was aligned to Mtb reference genome (Refseq accession: NC_000962.3) using BWA-MEM (v0.7.17). The alignment was sorted using samtools (v0.3.1) and the median fragment length was determined using bamPEFragmentSize (v3.5.4). It was used for calculating the GC bias frequencies using computeGCBias (v3.5.4). Finally, the GC error was corrected using correctGCBias (v3.5.4). The reads in fastq format were *de novo* assembled using Megahit (v1.2.9). The assemblies were used for drug resistance (DR) and lineage profiling using TB-profiler (v6.2.1) (6). The assemblies were used as input for synthesising reads using wgsim (v0.3.1) and mapped on the Mtb reference genome using the BWA-MEM algorithm.

### Matrix preparation and analysis

The aligned reads were counted using Bedtools coverage (v2.30.0). The read count matrix and grouping information synthesised using the output of the TB profiler were used to determine the input of gene copy number differences. The read count matrix was normalized using Variance Stabilizing Transform (VST). VST, unsupervised hierarchical clustering, and read count ratio analysis were conducted on the iDEP (v2.0) platform (42). Sva-Combat (v3.2.0) tool was employed for batch correction during multigroup analysis. ShinyGO v0.80 was used for gene ontology enrichment analyses. MS Excel 2016 home edition, GraphPad Prism (v8.4.3) and TB-tools (v1.112) were used for data representation. All data analysis was conducted on a server running an Intel Xeon E5-2630 v4 CPU with 135 GB of random-access memory.

### Approaches for relative gene copy number difference determination

Approach-I: Alignment of raw reads, sorting, read counting, and Variance stabilizing transformation (VST). Approach-II: Alignment of raw reads, sorting, GC bias correction, read counting and VST. Approach-III: Alignment of raw reads, sorting, GC bias correction, read counting, VST, and batch correction with sva. Approach-IV: De novo assembly, read simulation, alignment of raw reads, sorting.

### Statistical Analysis

Genes with Log_2_(fold change of reads aligning to genes) > ±1 and BH adjusted P value < 0.01 in read counts determined using DeSeq2 were considered to have significantly deregulated gene copy numbers between the groups analyzed.

## Declarations

### Consent for publication

Not applicable.

### Availability of data and materials

Publicly available WGS sequencing datasets in the European Nucleotide Archive were used in the present study (Supplementary Table 1).

### Authors’ contributions

NB and RKN conceptualized the study; NB performed formal analysis and prepared the first draft of the manuscript; AB mined literature databases for Mtb-specific BioProjects; AG contributed to data analysis using R tools. RKN provided administrative support and supervision and finalized the manuscript.

## Acknowledgements

Nidhi, Mothe Sravya, and Suchitra Jena from the Translational Health Group, ICGEB, New Delhi, are highly acknowledged for providing feedback and proofreading of the manuscript.

## Author’s information

Nikhil Bhalla (Ph.D.), Senior Research Fellow, Translational Health Group, ICGEB, New Delhi

Anil Behera, Junior Research Fellow, Translational Health Group, ICGEB, New Delhi

Ashish Gupta, Senior Research Fellow, Translational Health Group, ICGEB, New Delhi

Ranjan Kumar Nanda (Ph.D.), Group Leader, Translational Health Group, ICGEB, New Delhi

