## Supplementary Figure 1, Supplementary Table 1 for "Copy number variation analysis of 9,482 *Mycobacterium tuberculosis* isolates identifies lineage-specific molecular determinants"

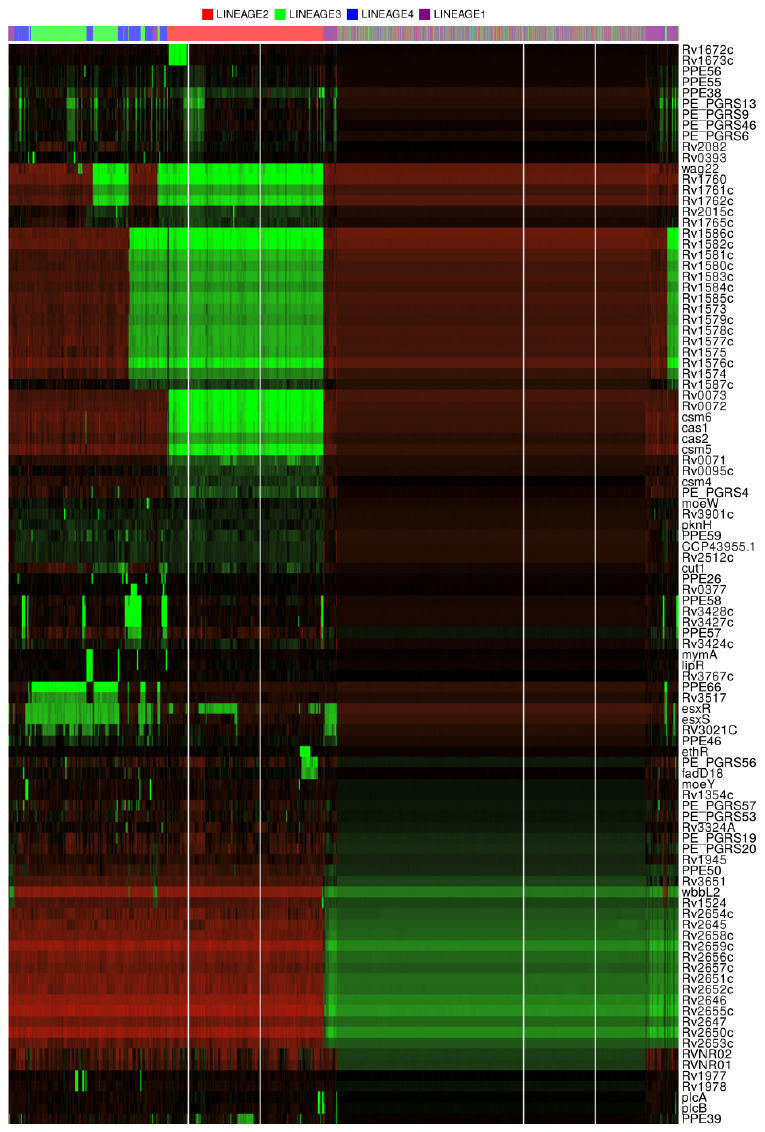

**Supplementary Figure 1: Unsupervised hierarchical clustering showed tandem genes with gene copy number differences between Mtb lineages.** The raw read counts of GC-corrected BAM files were subjected to VST, and the top 100 genes were used for heatmap generation, which showed genes with tandem occurrence on the Mtb genome. The heatmap subjected both columns (samples) and rows (gene IDs) to unsupervised hierarchical clustering. The scale is green-black-red for low-medium-high gene copy numbers.

**Supplementary Table 1:** Details of sources used for acquiring the Mtb whole genome sequencing (WGS) data for gene copy number difference analysis. *WGS raw data was not found in the European Nucleotide Archive (ENA). **The complete study references have been appended in the supplementary file.

| BioProject Accession | Country | Downloaded datasets | Filtered datasets | PMID | Study Reference** |
| --- | --- | --- | --- | --- | --- |
| PRJNA605605 | Bangladesh | 16 | - | 32612587 | 1 |
| PRJNA494931 | Brazil | 3 | - | 30514494 | 2 |
| PRJNA630228 |  | 69 | 42 | 33533871 | 3 |
| PRJNA756844 | Cambodia | 80 | 80 | 35562174 | 4 |
| PRJNA268900 | China | 606 | 533 | 30082724 | 5 |
| PRJNA436454 |  | 423 | 357 | 37271100 | 6 |
| PRJNA804712 |  | 43 | 42 | 35624432 | 7 |
| PRJNA806507 |  | 182 | 117 | 35311541 | 8 |
| PRJEB56100 | India | 246 | 42 | 36815861 | 9 |
| PRJNA822663 |  | 23 | 23 | 35442078 | 10 |
| PRJNA358480 |  | 5 | 5 | 30197919 | 11 |
| PRJNA996640 |  | 242 | 210 | 37671895 | 12 |
| PRJNA717132 |  | 65 | 65 | 35205308 | 13 |
| PRJNA379070 |  | 200 | 136 | 30863380 | 14 |
| PRJNA613351 |  | 82 | 77 | 33741489 | 15 |
| PRJNA512047 |  | 10 | - | 31784670 | 16 |
| PRJEB34541 |  | 78 | 5 | 32279871 | 17 |
| PRJEB41116 |  | 4301 | 4148 | 35989319 | 18 |
| PRJNA492975 |  | 32 | - | 30938706 | 19 |
| PRJNA885615 |  | 24 | 16 | 36695572 | 20 |
| PRJNA575883 |  | 2257 | 1559 | 37168873 | 21 |
| PRJNA224116 | Indonesia | 7 | - | 34802420 | 22 |
| PRJNA566379 | Iran | 37 | 37 | 32046149 | 23 |
| PRJNA957554 | Israel | 142 | 129 | 37928179 | 24 |
| PRJNA633244 | Java, Indonesia | 30 | 21 | 32579102 | 25 |
| PRJEB29276 | Lebanon | 19 | 11 | 30594126 | 26 |
| PRJNA575111 | Malaysia | 10 | 8 | 34989616 | 27 |
| PRJNA751891 | Mexico | -* | - | 35909960 | 28 |
| PRJEB48275 | Mixed | 215 | 208 | 35907429 | 29 |
| PRJEB42771 |  | 69 | 65 | 35741753 | 30 |
| PRJEB10037 | Myanmar | 14 | 8 | 27530852 | 31 |
| PRJNA645523 |  | 109 | 81 | 33007895 | 32 |
| PRJNA977967 | Oman | 69 | 69 | 37768070 | 33 |
| PRJEB32684 | Pakistan | 81 | 79 | 31628383 | 34 |
| PRJEB7798 |  | 42 | 40 | 25719196 | 35 |
| PRJEB43284 |  | 51 | 48 | 34244539 | 36 |
| PRJEB25972 |  | 641 | 632 | 34244055 | 37 |
| PRJNA629298, PRJNA629388 |  | 2 | 2 | 33862292 | 38 |
| PRJEB51057 | S. Korea | 71 | 58 | 37193005 | 39 |
| PRJNA879962 | Taiwan | 199 | 167 | 36781938 | 40 |
| PRJNA390471 | Thailand | 400 | 362 | 30981926 | 41 |
| PRJNA523980 |  | 71 | - | 36138798 | 42 |
| PRJNA972846 |  | 38 | - | 37992075 | 43 |
